# SINEultaneous profiling of epigenetic heterogeneity and transcriptome in single cells

**DOI:** 10.1101/2021.03.25.436351

**Authors:** Kooper V Hunt, Sean M Burnard, Ellise A Roper, Danielle R Bond, Matthew D Dun, Nicole M Verrills, Anoop K Enjeti, Heather J Lee

**Affiliations:** School of Biomedical Sciences and Pharmacy, College of Health Medicine and Wellbeing, University of Newcastle, Callaghan NSW Australia; Hunter Medical Research Institute, New Lambton Heights NSW Australia; NSW Health Pathology North, New Lambton Heights NSW Australia; Calvary Mater Newcastle, Waratah NSW Australia

## Abstract

Global changes in DNA methylation are observed in developmental and disease contexts, and singlecell analyses are highlighting the heterogeneous regulation of these processes. However, technical challenges associated with single-cell analysis of DNA methylation limit these studies. We present single-cell transposable element methylation sequencing (scTEM-seq) for cost-effective estimation of global DNA methylation levels. By targeting high-copy LINE-1 and SINE Alu elements, we achieve amplicon bisulphite sequencing with thousands of loci covered in each library. Parallel transcriptome analysis is also performed to link global DNA methylation heterogeneity with gene expression. We apply scTEM-seq to KG1a acute myeloid leukaemia (AML) cells, and primary AML cells. Decitabine treatment of KG1a cells induces global DNA methylation heterogeneity associated with altered expression of immune process genes. We also compare global levels of DNA methylation to expression of transposable elements and find a predominance of negative correlations in both the KG1a and patient cells. Finally, we observe co-ordinated upregulation of many transposable elements in a sub-set of decitabine treated cells. By linking global DNA methylation heterogeneity with transcription, scTEM-seq will refine our understanding of epigenetic regulation in cancer and beyond.

## INTRODUCTION

Single-cell analysis of DNA methylation has revealed epigenetic heterogeneity in development and disease, and parallel transcriptomic analyses are allowing this heterogeneity to be understood in the context of genomic regulation [1, 2]. For example, single-cell analysis of DNA methylation, chromatin accessibility and gene expression has demonstrated that active epigenetic remodelling is required for endoderm and mesoderm specification during gastrulation [3]. In contrast, the ectoderm lineage is epigenetically primed in the epiblast and serves as a default differentiation pathway. Similar analyses have been applied to colorectal cancer revealing relationships between somatic copy number alterations, DNA methylation and gene expression [4]. While genetic sub-lineages were found to have distinct epigenetic profiles, comparison between primary and metastatic sites suggested that epigenetic reprogramming was not essential for tumour dissemination.

The studies described above demonstrate the power of linking DNA methylation heterogeneity with genetic and transcriptional heterogeneity. However, technical challenges continue to limit the implementation of single-cell DNA methylation analyses. Most methods rely on bisulphite conversion to distinguish methylated from unmodified cytosines. This chemistry provides single-nucleotide resolution but is incompatible with available high-throughput droplet barcoding technologies. Thus, single-cell analysis of DNA methylation is currently limited to low-throughput multi-well plate assays that are relatively high cost. Furthermore, genome-wide bisulphite sequencing (BS-seq) requires ten times as many reads as RNA sequencing (RNA-seq), meaning that studies on thousands of cells are usually cost-prohibitive. Finally, the sparse data obtained from single-cell BS-seq (scBS-seq) and single-cell RNA-seq (scRNA-seq) libraries poses a major challenge to multi-omic studies hoping to identify individual loci where DNA methylation correlates with gene expression. In the study of colorectal cancer mentioned above [4], promoters with differential DNA methylation between primary tumour and metastatic sites were identified, but no correlations to expression of the associated genes were reported. Indeed, the most exciting findings from this study were related to global changes in DNA methylation, as opposed to locus-specific effects.

We reasoned that assessment of global DNA methylation in single cells would be a useful alternative to genome-wide analyses in contexts such as embryonic development and cancer, and reckoned that transposable element (TE) methylation could be exploited for this purpose. TEs are conserved DNA sequences capable of replicating and inserting into new positions in the genome. Discovered by Barbara McClintock in 1950 [5], TEs are estimated to make up around half of the human genome [6]. Poly-A retrotransposons Long Interspersed Element 1 (LINE-1) and Short Interspersed Element Alu (SINE Alu) account for almost 25% of the genome and are some of the only active or ‘hot’ transposable elements still capable of transposing in our genome [7, 8]. Active retrotransposition causes genome instability, and because of this mutagenic potential, TEs are epigenetically silenced by high DNA methylation levels in internal promoters. Consequently, global changes in DNA methylation are correlated to changes in TE methylation in early embryonic development [9], primordial germ cell development [10], induced pluripotent stem cell (iPSC) reprogramming [11] and cancer [12]. Indeed, even in the single-cell analysis of colorectal cancer discussed above, lineagespecific global DNA hypomethylation was associated with an over-representation of TE sequences (LTRs, LINEs) [4].

These observations justify the use of TEs as surrogate measures for global DNA methylation levels, and LINE-1 and SINE Alu elements are common targets for bisulphite conversion-based analysis [13, 14]. Here we adapt this approach for cost-effective analysis of global DNA methylation levels in a method called single-cell transposable element (TE) methylation sequencing (scTEM-seq). To achieve this we perform targeted amplification of bisulphite converted SINE Alu and LINE-1 sequences. We apply scTEM-seq in acute myeloid leukaemia (AML) cells and detected global DNA methylation heterogeneity following treatment with the hypomethylating agent decitabine (DAC). Parallel analysis of gene expression in the same single cells identified links to viral response pathways correlated after DNA methylation loss.

## MATERIAL AND METHODS

### Cell Lines and Patient Samples

KG1a cells were cultured in Iscove’s Modified Dulbecco’s Medium (IMDM) (Sigma-Aldrich, catalog # I3390) with 10% fetal bovine serum (FBS). Cell lines were treated with 100nM 5-aza-2’-deoxycitidine (decitabine, DAC) every 24 hours (0, 24 and 48 hours) and harvested at 72 hours.

The AML patient included in this study (AML01) was recruited at diagnosis through the Calvary Mater Newcastle Hospital, with written informed consent. Studies were approved by the human ethics committees of the Hunter New England Area Health service, and the University of Newcastle. The patient was a 60-year-old male, diagnosed with secondary AML following chronic myelomonocytic leukaemia. Clinical assessment revealed a complex karyotype including an isochromosome 17q, and mutations in the *ASXL1, SETBP1* and *SRSF2* genes. Enriched mononuclear cells were purified from peripheral blood using Lymphoprep density gradient medium (StemCell, catalog # 7851) and SepMate tubes (StemCell, catalog # 85450), and cryopreserved.

### Cell Sorting

KG1a cells were stained using the PE Annexin V Apoptosis Detection Kit (BD Life Science, catalog # 559763). Live cells (Annexin V^-^/7-AAD^-^) were sorted into individual wells of a 96 well plate containing lysis buffer 2.5μL RLT Plus Lysis Buffer (QIAGEN, catalog # 1053393) with 1U/μL SUPERase-In (ThermoFisher, catalog # AM2696). Before sorting, bulk samples of 1000000 cells were collected from both the untreated and treated populations for comparison with single cells.

Cryopreserved primary human cells were resuspended in thawing media (IMDM, 20% FBS), washed twice and resuspended. The cells were then rested for 1h at 37°C before preparation for flow cytometry. Cells (1×10^6^/100ul) were stained with 1.5μg/mL propidium iodide (PI, Sigma-Aldrich, P1304MP), 1:20 CD45-PECy7 (2D1, Life Technologies, catalog #25-9459-42), 1:20 CD33-FITC (WM-53, Life Technologies, catalog # 11-0338-42) and 1:20 CD19-BV711 (SJ25C1, BD Biosciences, catalog # 563036). Single blasts (PI^-^/CD45^dim^) were collected in 2.5μL RLT Plus Lysis Buffer containing 1U/μL SUPERase-In in 96 well plates.

### Library Preparation

We utilised the G&T-seq protocol to separate genomic DNA and RNA from the single-cell samples [15]. Genomic DNA from each cell was purified and bisulphite conversion was performed as described [16], with minor modifications. Bisulphite conversion was carried out using the EZ-96 DNA Methylation-Direct MagPrep Kit (Integrated Sciences, catalog # D5054) with half volumes of the manufacturer’s instructions. Bisulphite converted DNA was eluted directly from MagBeads into PCR-mix, and amplification of TEs was performed with MagBeads still in the well. PCR cycling conditions used were 95°C for 5 min (1 cycle), 98°C for 20 sec, 53°C for 15 sec, 72°C for 1 min (35 cycles), and 72°C for 10min (1 cycle). PCR mix used 7.5μl 1x KAPA HiFi hotStart Uracil + ReadyMix (Millennium, catalog # ROC-07959079001) and 0.3μM primer mix. Primers were designed against SINE Alu and LINE-1 consensus sequences (Supplementary Figure S2A) and are contained in Supplementary Tables S1 and S2. After amplification, plates of 96 single cell libraries purified using a 1.2x volume of AMPure XP beads (Beckman Coulter, catalog # A63881). All libraries were then quantified using the Qubit dsDNA HS kit (Life Technologies), normalised and pooled to a single tube. Pools were then added to 0.8μM NEBNext dual index oligo sets (Genesearch, catalog # E7780S) and 14.5μl 1x KAPA HiFi HotStart ReadyMix (Millennium, catalog # ROC-07958935001) for indexing and adaptor addition. PCR cycling conditions used were 98°C for 45 sec (1 cycle), 98°C for 15 sec, 65°C for 30 sec, 72°C for 30 sec (5 cycles), and 72°C for 5 min (1 cycle). Pools were then purified using 0.9x volume of Ampure XP beads, normalised and combined for sequencing. Matched scRNA-seq libraries were prepared as described [16].

A post-bisulphite adaptor tagging (PBAT) approach [17] was used to prepare genome-wide sequencing libraries from bulk samples. Libraries were prepared as described [18], with minor modifications. The 6NR adaptor 2 oligo used during second strand synthesis was modified (5’-CAGACGTGTGCTCTTCCGATCTNNNNNN-3’) to be compatible with NEBNext dual index oligos that were used for library amplification.

### Sequencing

Sequencing of bisulphite reads was performed using the Illumina MiSeq platform. Low read depth is required, so for data in this paper sequencing kits with only 4 million reads were used for 192 cells. Library loading concentrations of 8-10pM were used with a 1% PhiX spike-in. We achieved on average 23000 read pairs per sample.

scRNA-seq Libraries were sequenced using the NextSeq platform with a loading concentration of 1.5pM and a 1% PhiX spike-in. We excluded all cells with alignment rates under 80%. With approximately 1000000 reads per cell, we measured between 6300 and 15000 genes in all of our single cell KG1a scRNA-seq libraries (Supplementary Table S5). Gene numbers measured in AML01 cells were more modest, with between 2800 and 5200 genes in cells passing quality control (Supplementary Table S6).

PBAT libraries were sequenced using the MiSeq platform. These libraries were prepared with the intention of measuring global DNA methylation levels and as such were also sequenced with low read depth (~100000 reads per bulk sample).

### Data processing and analysis (scTEM-seq)

After initial demultiplexing of primary Illumina indexes, Cutadapt (v2.10) [19] was used to demultiplex pools based on custom secondary indexes (Supplementary Table S1). Commands –g and -G were used to pass named forward and reverse index lists as a .fasta file to Cutadapt. Bisulphite reads were trimmed using Trim Galore (v0.6.5) [20]. 10bp was trimmed from both the 5’ and 3’ ends to remove oligo sequences from reads. Reads were mapped to Bowtie2 (v 2.4.1) [21] indexed human genome (GRCh38) using Bismark (v0.22.3) in non-directional and paired-end mode [22]. The methylation extraction module from Bismark was then used to produce coverage files (.cov) for methylation analysis. We excluded cells with coverage of less than 1000 annotated TE sites. Downstream analysis was performed using Seqmonk [23].

### Data processing and analysis (PBAT)

PBAT libraries were trimmed using Trim Galore (v0.6.5) to remove 9bp from the 5’ end of all reads. Reads were mapped using Bismark (v0.22.3) in non-directional and paired-end mode. Unmapped reads were re-aligned in single-end mode to account for chimeric reads seen in PBAT libraries [24]. After producing coverage files with Bismark methylation extraction module, paired and single end alignments for each sample were merged into a single file using the cat (concatenate) command. Downstream analysis was performed using Seqmonk.

### Data processing and analysis (scRNA-seq)

scRNA-seq data was trimmed using Trim Galore (v0.6.5), with default setting in paired-end mode. Hisat2 [25] (v2.1.0) and Samtools [26] (v1.1.0) were used to convert, map and align unique and ambiguous reads to the human reference genome build GRCh38 from raw fastq reads into bam format. TEtranscripts [27] was used to obtain raw gene and transposable element counts from the unique and ambiguously aligned reads using the GTF files for 1) TEs (http://labshare.cshl.edu/shares/mhammelllab/www-data/TEtranscripts/TE_GTF/) and 2) genes (https://asia.ensembl.org/info/data/index.html; release 101 from the FTP server) in GRCh38 ensembl format. TEtranscripts was run in a Conda [28] environment setup with Python (v3.7.7), Pysam (v0.16.0.1), R-base (v4.0.3) and Bioconductor-Deseq2 (v1.28.0).

Analysis of correlation of gene expression and DNA methylation was performed using R [29]. We included transcripts with at least 2 reads in at least 10 cells. Read counts for scRNA-seq data were normalised per million reads for each sample. We then performed a simple Pearson’s correlation test between gene expression values and DNA methylation levels in each cell.

### Data processing and analysis (scBS-seq)

Analysis was performed on data published by Bian et al. [4], downloaded from European Nucleotide Archive (ENA) (study accession PRJNA382695).

First, raw reads were trimmed in single-end mode using Trim Galore (v0.6.5) with 9bp removed from the 5’ end of each read. Secondly, trimmed reads were mapped using Bismark (v0.22.3) in non-directional mode to a bowtie2 indexed human genome (GRCh38). Mapped reads were then deduplicated by Bismark with default settings. The methylation extraction module from Bismark was then performed on the deduplicated reads, with the resulting coverage files for read pairs merged into a single file. Methylation levels were then assessed using the GenomicRanges [30] package and in R.

## RESULTS

To investigate whether TEs might serve as surrogate measures for global DNA methylation levels in single cell data, we first interrogated genome-wide scBS-seq data from a colorectal cancer patient (CRC01) [4]. We observed very strong correlations between DNA methylation within TE annotations and genome-wide levels for both LINE-1 (*R*^2^=0.88, p<2.2^-16^) and SINE Alu (*R*^2^=0.91, p<2.2^-16^) families (Supplementary Figure S1A). Furthermore, TE methylation was sufficient to identify sub-clonal differences in global DNA methylation as previously reported (Supplementary Figure S1B) [4]. Thus, TE methylation in single cell data can highlight biologically interesting heterogeneity in cancer cells.

We modified the scBS-seq protocol and achieved amplification of SINE Alu and LINE-1 sequences following bisulphite conversion of single cell DNA samples. In initial experiments, SINE Alu primers delivered greater amplicon yield and library complexity than LINE-1 primers, consistent with the higher copy-number of SINE Alu elements (Figure 1A). A second generation of SINE Alu primer sequences was then designed to increase the number of scTEM-seq libraries that can be pooled for sequencing, and to ameliorate the technical challenges of sequencing low-diversity amplicon libraries (Supplementary Figure S2B, Supplementary Tables S1 and S2).

**Figure 1:**
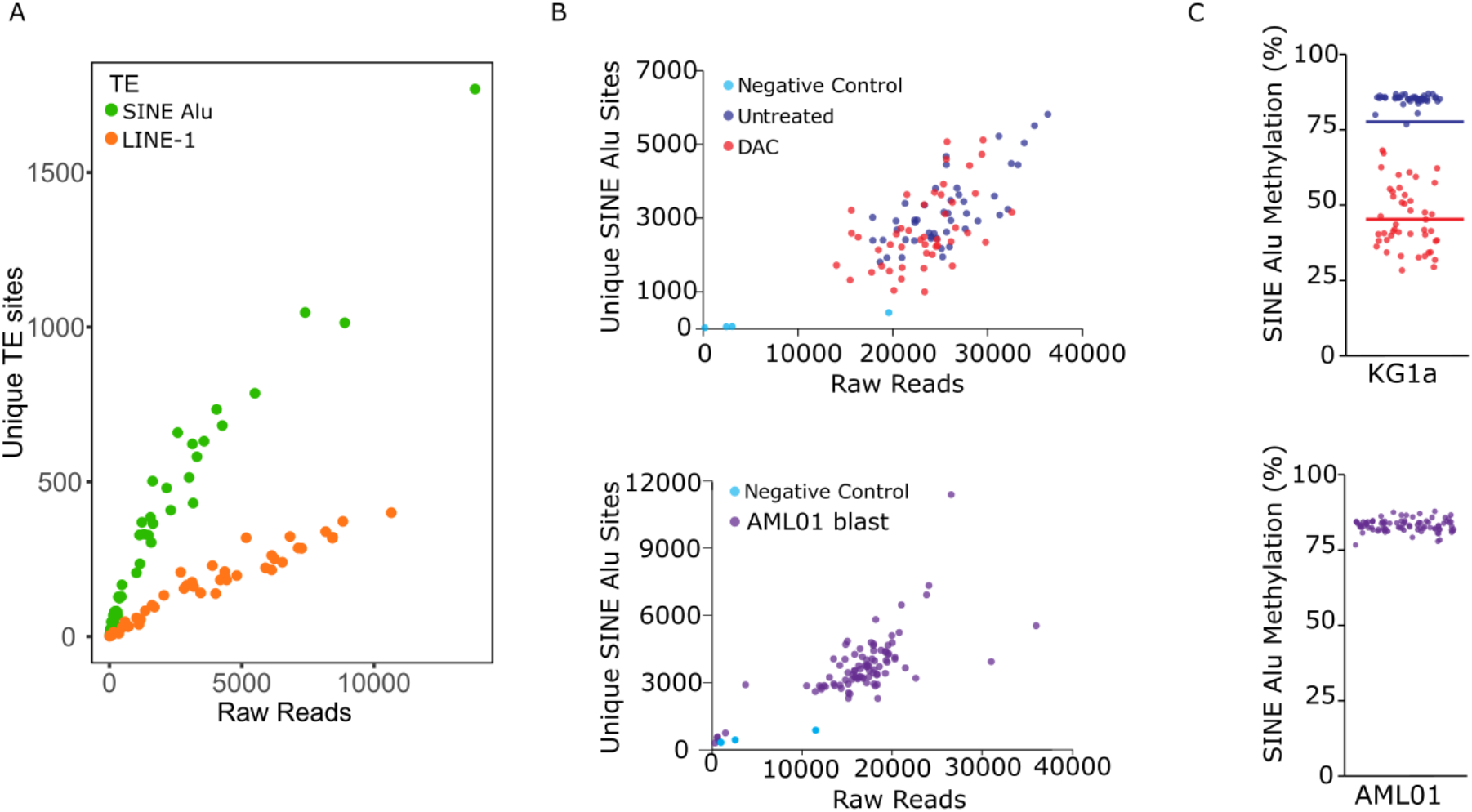
scTEM-seq accurately measures DNA methylation at TE sites. A) Comparison of unique TE sites measured against the number of uniquely mapped reads. Using scTEM-seq, greater coverage of SINE Alu was achieved per sequencing reads than LINE-1. B) Unique SINE Alu sites measured in KG1a cells and AML01 patient blasts compared to raw sequencing reads. C) DNA methylation levels as measured by scTEM-seq in KG1a cells with and without DAC treatment and untreated AML01 patient blasts. Coloured lines show average whole genome DNA methylation levels for each treatment group measured by post-bisulphite adapter tagging (PBAT). DAC treated KG1a cells show a heterogeneous loss of DNA methylation.

We applied our optimised scTEM-seq protocol to KG1a AML cells treated with and without 100nM DAC for 72h. Negative controls were also included to monitor DNA contamination in reagents. Average amplicon yield from single cell samples was 16.10ng/μl (4.08 SD), compared to 1.12ng/μl (1.03 SD) in negative controls (Supplementary Figure S3). scTEM-seq libraries achieved unique alignment rates of 67.23% (5.11 SD) (Supplementary Figure S3), and efficient bisulphite conversion was confirmed by very low non-CpG methylation rates (DNA methylation in CHG trinucleotide contexts was 0.67%, 0.2 SD) (Supplementary Table S3). In single cell samples, information was recovered from 1000-6000 unique SINE Alu annotations in the reference genome, despite low sequencing depth (14000-37000 raw reads) (Figure 1B). Untreated cells had uniformly high levels of DNA methylation in SINE Alu elements, with an average of 85.4% (1.65 SD). DAC treatment resulted in heterogeneous loss of DNA methylation with levels ranging from 29% to 69%, with an average of 41.86% (10.46 SD) (Figure 1C). We compared our scTEM-seq results to average genome-wide methylation levels from the same populations of KG1a cells using a PBAT approach. The bulk DNA methylation measurements were similar to scTEM-seq estimates at 43.87% and 78.58% in DAC treated and untreated cells, respectively (Figure 1C). This demonstrates that targeted transposable element sequencing can be used to estimate global DNA methylation levels in single cells.

To test scTEM-seq in primary human cells, we applied our analysis on sorted blasts from an AML patient. Amplicon yield, alignment rates and bisulphite conversion were comparable to KG1a samples, and 88 of 92 libraries passed quality control with representation of >1000 unique SINE Alu elements (Supplementary Table S4). This patient had not received hypomethylating agent therapy, and DNA methylation at SINE Alu elements was consistently high in these cells (84.74%, 2.15 SD) (Figure 1C).

Prior separation of gDNA and RNA from each cell allowed us to prepare matched scRNA-seq libraries for paired analysis of DNA methylation levels and gene expression. In our DAC treated KG1a data, 60 genes had a correlation after multiple testing correction with the majority (43) being positive associations (Figure 2A). For example, interferon induced protein *IFI44L* was down-regulated in cells with lower SINE Alu methylation (*R* = 0.24, p = 3.8e^-05^), whereas major histocompatibility complex (MHC) I component *HLA-A* was up-regulated in cells with lower SINE Alu methylation (*R*^2^ = 0.14, p = 2.0e^-04^) (Figure 2B). Gene ontology analysis on all genes with a significant correlation to DNA methylation (FDR<0.05) revealed several over-represented pathways, including terms related to immune processes (‘leukocyte mediated immune response’) and viral processes (‘biological process involved in interspecies interaction between organisms’) (Figure 2C).

**Figure 2:**
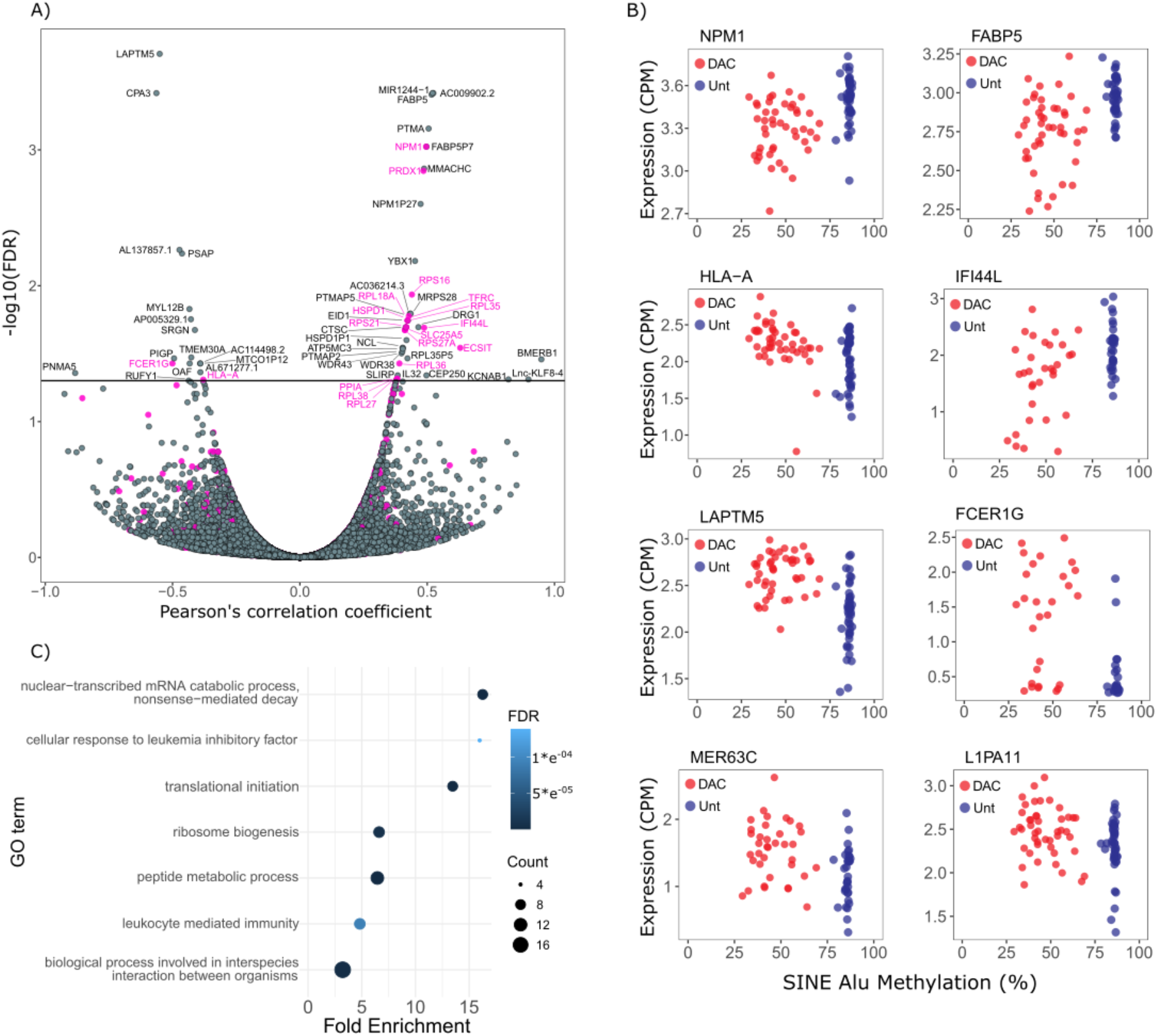
DNA methylation correlates with expression of immune genes. A) Volcano plot showing Pearson’s correlation between DNA methylation levels and gene expression in the KG1a dataset. B) Gene ontology (Panther) results for statistically overrepresented biological pathways in all genes with expression correlated to DNA methylation (FDR<0.05). For related terms, only the pathway with the highest number of correlated genes is displayed for simplicity. Genes included in ‘biological process involved in interspecies interaction between organisms’ were also annotated in a related ‘viral process’ term, and are highlighted in purple in (A). C) Select examples showing expression levels of an individual gene (in CPM or counts per million) and DNA methylation levels in our treated and untreated KG1a cells. Examples include 6 genes (*NPM1, FABP5, HLA-A, IFI44L, LAPTM5* and *SRGN*) and 2 TEs (MER63C and L1PA11).

Hypomethylating agents like DAC have been shown to act through a ‘viral mimicry’ process whereby loss of DNA methylation induces transcription of transposable elements (e.g. endogenous retroviruses, LINEs and SINEs) and a subsequent type 1 interferon response in effected cells [31–33]. To test whether scTEM-seq could detect expression of these TEs in single cells, we examined the abundance of TE sequences in transcripts from DAC treated and untreated KG1a cells. DAC treated cells showed a clear increase in TE expression levels compared to untreated cells (Supplementary Figure S4A). Although no TEs had significant correlation between DNA methylation and gene expression after multiple testing correction, we saw a trend towards negative correlation in both of our datasets (Supplementary Figure S4B). To further investigate TE expression patterns we performed clustering analysis of TE families that were differentially expressed after DAC treatment (Figure 3, Supplementary Figure S5). In our KG1a cells, we observed a subgroup of mostly DAC treated cells with co-ordinated up-regulation of many TEs, especially LINE-1 and SINE Alu families (Figure 3). Interestingly, this subgroup of cells could not be distinguished from other DAC treated cells based on global DNA methylation alone (44.75% (9.34 SD) vs 48.02% (11.16 SD), respectively), suggesting that other factors must regulate TE expression in the absence of DNA methylation.

**Figure 3:**
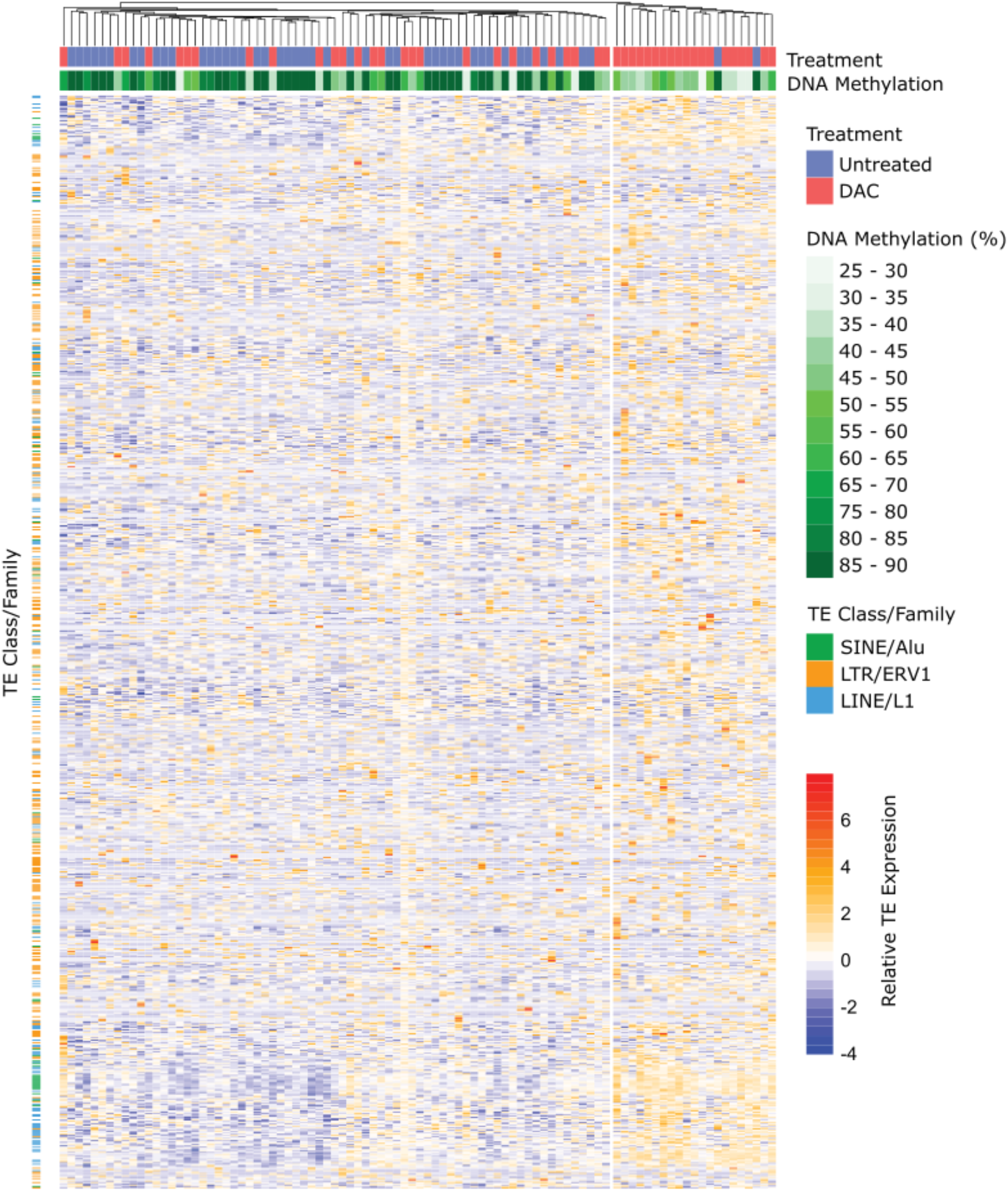
Coordinated up-regulation of TE transcription is observed in a subset of DAC treated KG1a cells. TEs with differential expression between untreated (blue) and DAC treated (red) KG1a cells (adjusted p<0.05) were grouped by family and normalised read counts were computed by variance stabilisation transformation (vst) in DESeq2. The heatmap shows relative TE expression levels with z-score scaling applied to each row, and cells clustered by Euclidean distance. Global DNA methylation percentages for each cell are indicated (green scale at top) and selected TE families are highlighted (left). A sub-cluster of mostly DAC treated cells (red) have low DNA methylation and high expression of TEs.

## DISCUSSION

TEs have been widely targeted for surrogate measures of global DNA methylation. We have adapted this approach to single cells, developing a cost-effective alternative to genome-wide techniques [34–39]. While other studies have amplified loci of interest in bisulphite converted DNA from single cells [40–44], ours is the first to target TEs. Since SINE Alu and LINE-1 elements are present at high copy number, our method requires far fewer cycles of amplification limiting the impact of PCR biases and errors. We demonstrate that methylation of SINE Alu elements in single cells compares well to global DNA methylation levels using *in silico* analysis of published data (Supplementary Figure S1) and by comparing scTEM-seq to matched bulk PBAT libraries (Figure 2C). In each case, SINE Alu methylation is shown to slightly over-estimate global DNA methylation levels (e.g. by 6.8% for untreated KG1a cells) (Figure 2C), which can be explained by the well-characterised enrichment of TEs in hypermethylated regions of the genome [45].

scTEM-seq has several advantages over comparable genome-wide methods such as single-cell bisulphite sequencing (scBS-seq) [35]. scTEM-seq libraries are prepared using sequence-specific primers, rather than random-priming oligos, leading to reduced oligo contamination and improved alignment rates. Indeed, the unique alignment rates for scTEM-seq libraries are surprisingly high considering that repetitive loci are inherently difficult to map in the reference genome. Improved alignment rates confer a cost saving by reducing wastage from sequencing runs; however, an even greater advantage is obtained by reducing the sequencing demand. Whereas scBS-seq libraries require ~20 million reads per cell to obtain genome-wide information, scTEM-seq libraries can provide a global estimate of DNA methylation from ~20 thousand reads. Thus, the sequencing cost is 3 orders of magnitude lower for scTEM-seq libraries. Obviously, this reduced cost comes with a considerable loss of information. However, locus-specific analysis of DNA methylation is also difficult in genomewide libraries, due to the low coverage obtained in each cell (e.g. 10-40% of the genome). Like scBS-seq and other plate-based methods, scTEM-seq is compatible with parallel analysis of gene expression in the same single cell. This allows epigenetic heterogeneity to be linked to transcriptional output. Thus, scTEM-seq will increase the scale of single cell studies in biological contexts where global changes in DNA methylation are of interest.

In this study, DAC treatment of KG1a AML cells led to heterogeneous loss of DNA methylation and altered expression of many genes (Figure 2). For example, *HLA-A* and *FCER1G* were negatively correlated to DNA methylation only 72 hours after initial treatment, possibly signifying an early immune response in cells that have lost DNA methylation. Furthermore, we were able to link epigenetic heterogeneity to expression of TEs, suggesting that variable activation of viral mimicry pathways could influence treatment response. We identified a subgroup of DAC treated KG1a cells with co-ordinated up-regulation of many TE families. This group of cells could not be distinguished based on DNA methylation levels alone, suggesting that loss of methylation is insufficient for activation of viral mimicry. In cells that do not up-regulate TEs, other epigenetic processes may substitute for the suppressive effects of DNA methylation, or transcriptional activators required for TE expression may be absent. A major limitation for the clinical use of hypomethylating agents is the variability in patient response. Although azacitidine has been shown to improve survival compared to conventional care, a large proportion of patients receive little or no benefit [46]. Analysis of methylation levels, measured in bulk tumour samples, has not been able to predict patient response to azacitidine [47]. Expression of subsets of evolutionarily young TEs however can enhance the viral mimicry response and correlates with improved prognosis [32]. Using scTEM-seq, we can take these studies a step further and explore how heterogeneity of DNA methylation and expression of TE subtypes within a tumour contribute to patient prognosis.

We also applied scTEM-seq to primary patient blasts, revealing homogeneous levels of DNA methylation. We did not identify correlations between DNA methylation levels and gene expression in this small set of cells (data not shown). However, we did note a bias toward increased TE expression in cells with lower DNA methylation levels. This is consistent with previous observations that DNA methylation proximal to TE sites correlates with their expression across different cancer types [48].

Thus, even in relatively homogeneous populations of cancer cells, variation in TE methylation may lead to intra-tumoural heterogeneity in TE expression and viral mimicry pathways. Future studies will apply scTEM-seq to many cells from numerous patients to test whether this observation represents a general phenomenon in cancer.

scTEM-seq will also be applicable to other aspects of cancer biology. Common mutations in epigenetic regulators (e.g. *DNMT3A, TET2, IDH1/2*) are known to cause widespread dysregulation of the cancer methylome, but their impact on epigenetic variability is unclear. scTEM-seq could be used to assess how altered turnover of cytosine modifications impacts intra-tumoural heterogeneity in cancers with these mutations. Future studies may also infer copy number alterations and specific gene variants from analysis of parallel single-cell transcriptome data [49, 50]. This would allow global changes in DNA methylation to be compared between genetic sub-clones [4]. scTEM-seq is also relevant to several contexts in stem cell and developmental biology. iPSC reprogramming is a heterogeneous process in which global epigenetic remodelling accompanies reactivation of pluripotency networks [51, 52]. Variable DNA methylation in iPSCs raises concerns regarding their safety in clinical regenerative medicine since incorrect reprogramming could lead to cancerous growth [53]. Thus, scTEM-seq may be a useful tool to understand the heterogeneity and assess the quality of iPSCs. As well as reporting global changes in DNA methylation, scTEM-seq may be applied to understand TE biology and activation during development [54]. TEs are also a source of regulatory elements that can become active to drive endogenous gene expression during development and oncogene expression in cancer [55–57]. Thus, scTEM-seq could be used to link TE methylation to expression networks even in the absence of global DNA methylation shifts. Of course, other species also have high-copy transposable element sequences and DNA methylation, so scTEM-seq could be adapted to many organisms. Ultimately, scTEM-seq will find applications in many aspects of medicine and biology. The reduced complexity and cost of this approach will also allow multi-dimensional single-cell analysis to be used more often and at scale.

## DATA AVAILABILITY

Raw sequencing data together with processed files (RNA counts and CpG methylation reports) are available from the authors upon reasonable request and will be made publicly available upon final publication.

## ACKNOWLEDGEMENT

We thank Dr Shuhui Bian (Peking University) for providing the genetic sub-lineages of CRC01 patient cells analysed in Supplementary Figure S1. We thank Dr Carlos Riveros (Hunter Medical Research Institute) for consultation regarding data processing pipelines related to this work, and Dr Shalin Naik (Walter and Eliza Hall Institute) for suggestions regarding primer design. We also thank Professor Geoffrey Faulkner and Dr Sandra Richardson (University of Queensland) for helpful discussions regarding transposable element biology, and Ms Nicole Cole (Hunter Medical Research Institute) for assistance with fluorescence activated cell sorting.

## SUPPLEMENTARY FIGURES

**Supplementary Figure S1:**
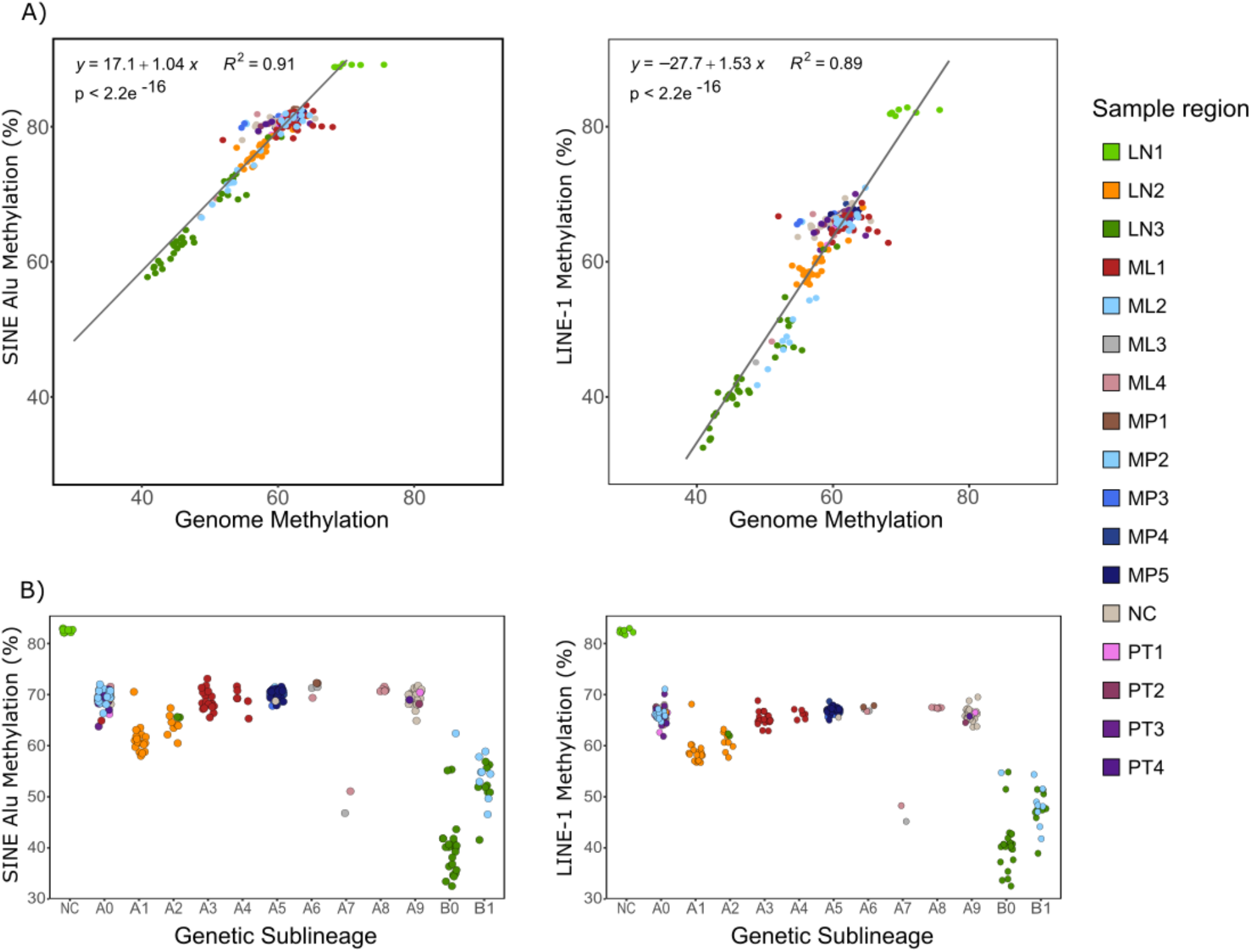
Transposable element DNA methylation accurately reflects global levels in single-cell data. Single-cell data from a colorectal cancer patient (CRC01) were obtained from Bian et al. 2018 (GSE97693). Bisulphite sequencing libraries were processed as described in methods and DNA methylation averages were calculated for all cytosines, or those within TE annotations. A) DNA methylation in TEs (SINE Alu and LINE-1) correlates with genome wide levels. R^2^ and p values calculated using Pearson’s correlation. B) Genetic sub-lineages were defined by copy number analysis in Bian, Hou et al. 2018. DNA methylation in TEs was sufficient to detect the global hypomethylation observed in sub-lineages B0 and B1. All plots are coloured by sampling regions. LN = lymph node metastasis, ML = liver metastasis, MP = post-treatment liver metastasis, NC = adjacent normal colon, PT = primary tumour.

**Supplementary Figure S2:**
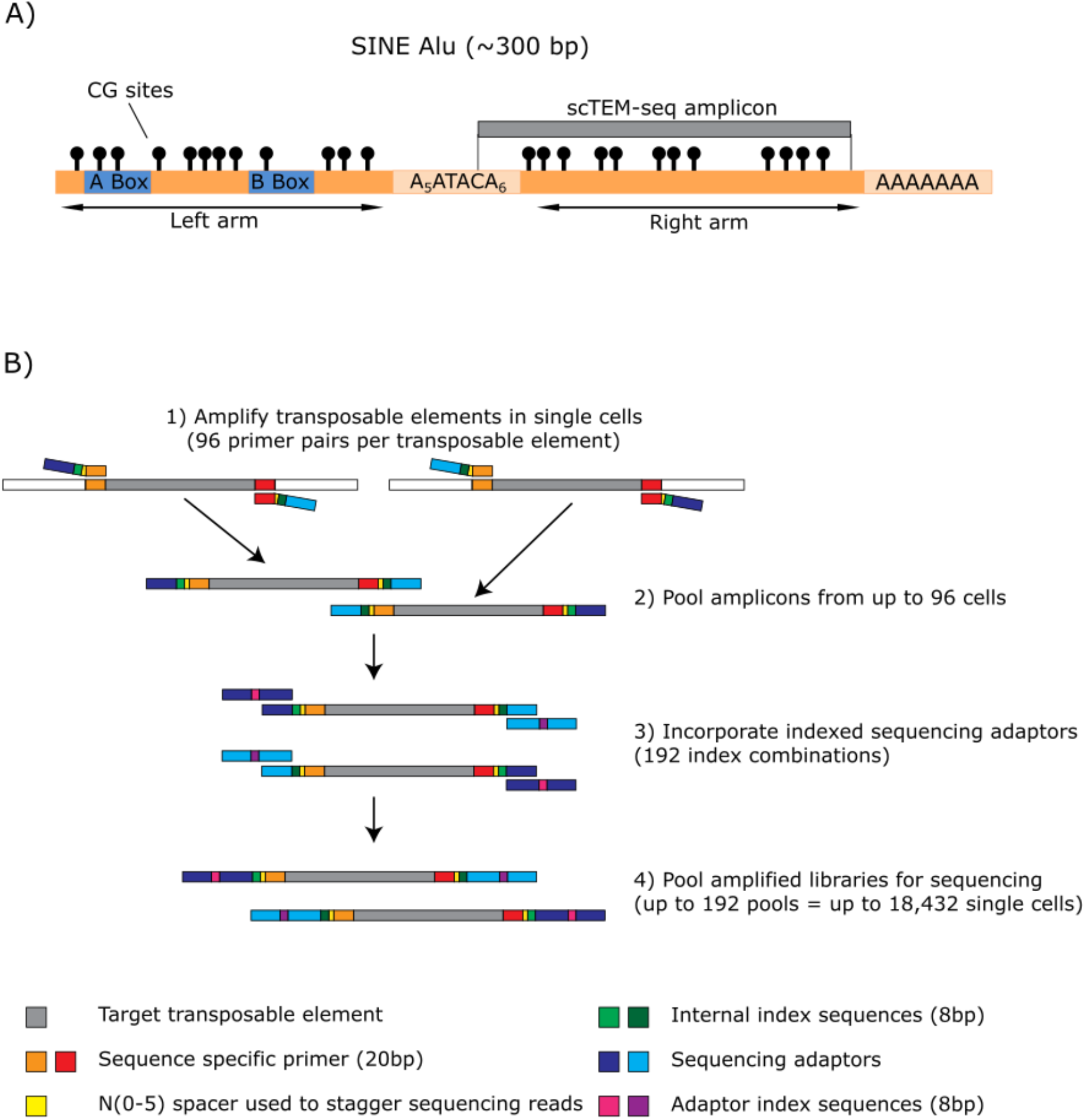
Amplicon and primer design for scTEM-seq. A) Location of SINE Alu amplicon in relation to the consensus sequence. B) scTEM-seq library preparation showing amplification and incorporation of index and adapter sequences. Sequencespecific amplification of SINE Alu elements is first performed using oligos carrying partial adaptor sequences. A spacer of 0-5 random nucleotides is included to artificially increase library complexity by phasing amplicon reads. Internal dual index sequences are also included so that 96 single cell libraries can be pooled before a second amplification is used to introduce full sequencing adaptors. The adaptors also carry dual index sequences, so that up to 192 pools of 96 single cells can be sequenced in parallel.

**Supplementary Figure S3:**
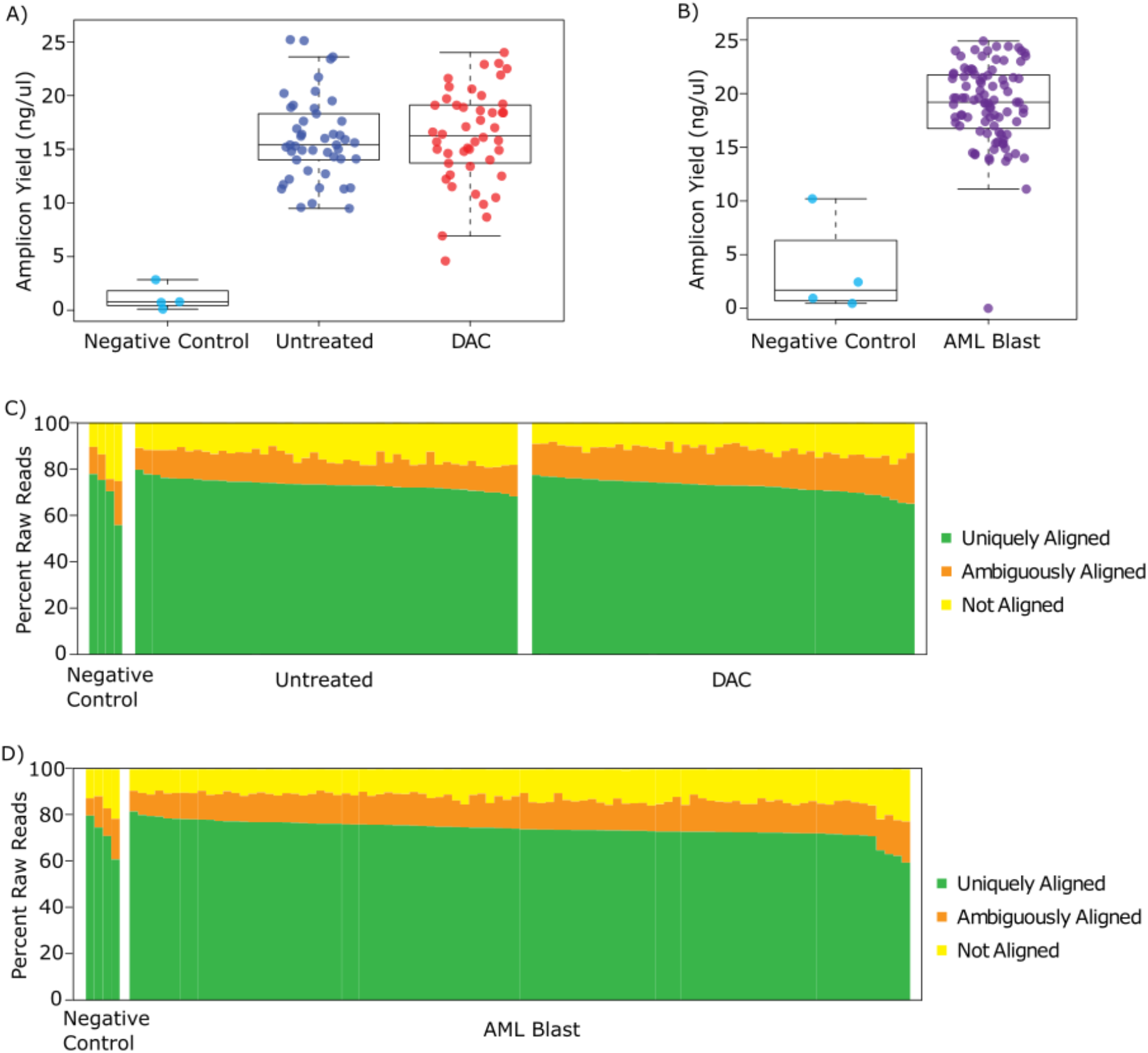
Quality control of scTEM-seq libraries. A) Amplicon yield and C) mapping efficiency in DAC treated and untreated KG1a cells and negative controls. B) Amplicon yield and D) mapping efficiency in untreated AML01 patient cells and negative controls.

**Supplementary Figure S4:**
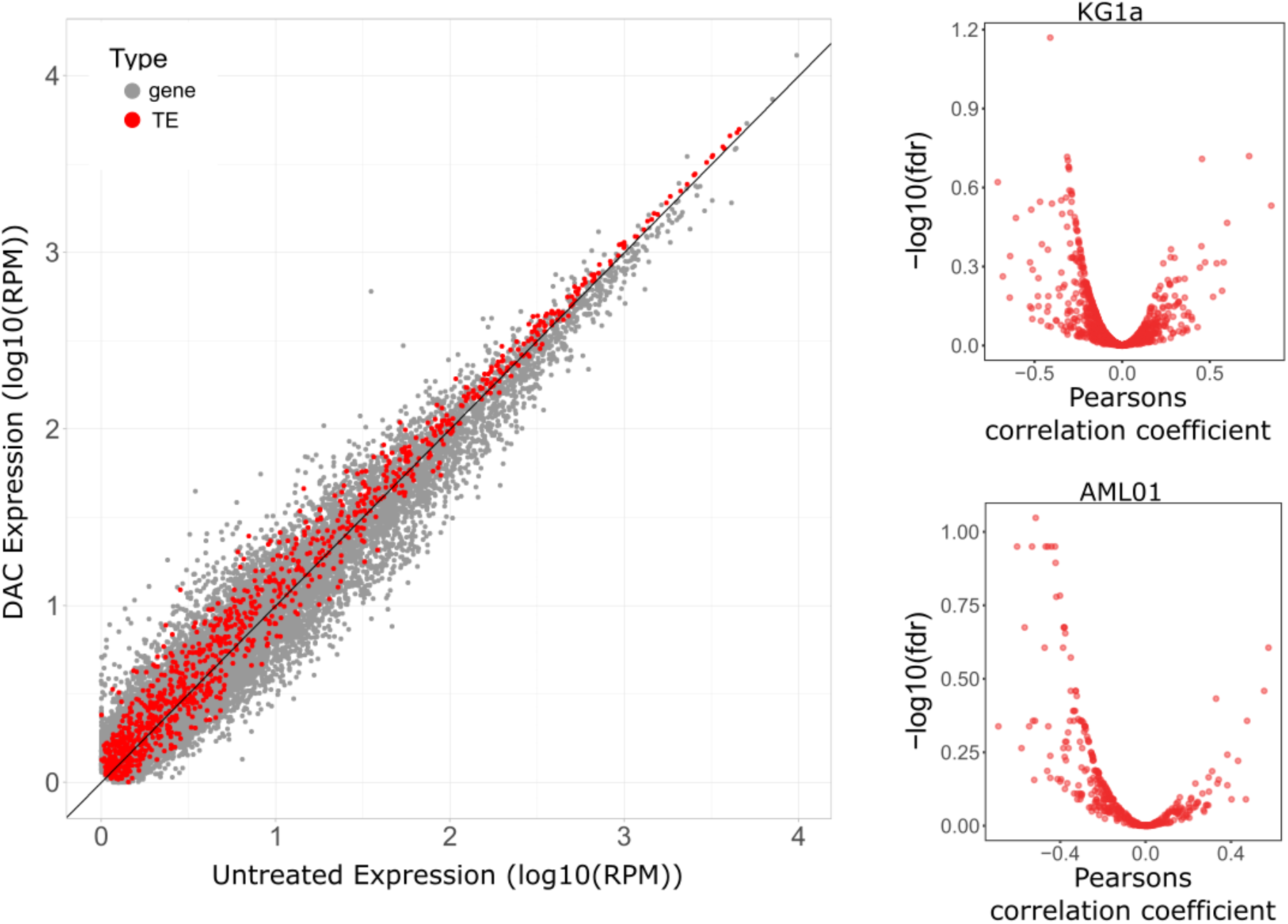
Transcription of TEs in single-cell data. A) Comparison of the average normalised expression values (log10(RPM)) for genes (grey dots) and TEs (red dots) in DAC treated vs untreated KG1a cells. An intersect line (y=x) is plotted with any gene or TE above the line representing a relative increase in average expression in DAC treated compared to untreated. B) Volcano plot showing Pearson’s correlation between DNA methylation levels and TE expression for KG1a and AML01 datasets.

**Supplementary Figure S5:**
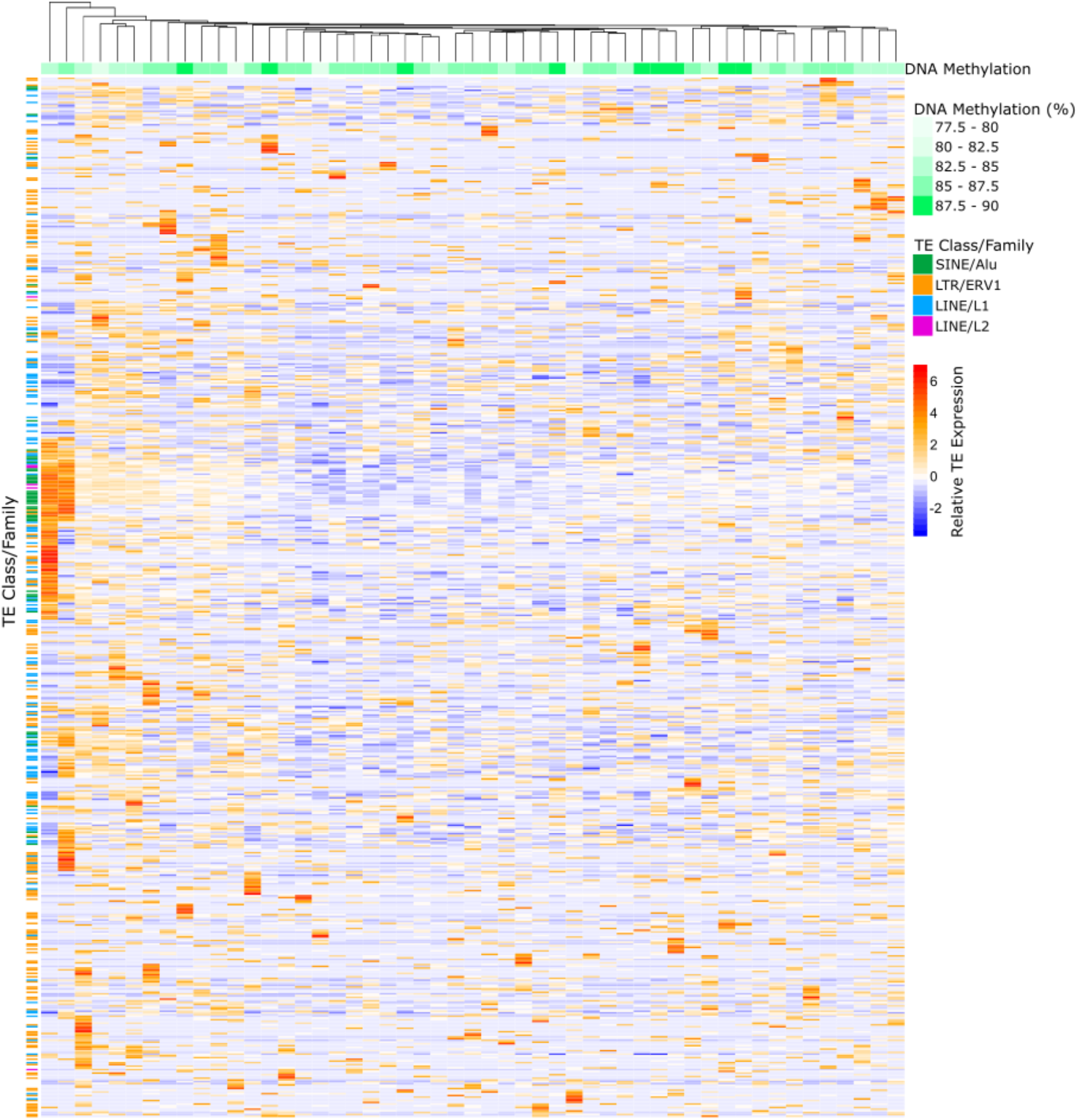
Heatmap of TE expression in AML01 blasts. TEs with differential expression between untreated and DAC treated KG1a cells (adjusted p<0.05) were grouped by family and normalised read counts were computed by variance stabilisation transformation (vst) in DESeq2. The heatmap shows relative TE expression levels in AML01 cells with z-score scaling applied to each row, and cells clustered by Euclidean distance. Global DNA methylation percentages for each cell are indicated (green scale at top) and selected TE families are highlighted (left).

## SUPPLEMENTARY TABLES

**Supplementary Table S1: scTEMseq primer sequences.**

**Supplementary Table S2: scTEMseq primer pairs (96 well plate layout).**

**Supplementary Table S3: Quality control of scTEM-seq libraries prepared from KG1a AML cells.**

**Supplementary Table S4: Quality control of scTEM-seq libraries prepared from primary AML cells.**

**Supplementary Table S5: Quality control of scRNA-seq libraries prepared from KG1a AML cells.**

**Supplementary Table S6: Quality control of scRNA-seq libraries prepared from primary AML cells.**

